# MicrobeAnnotator: a user-friendly, comprehensive microbial genome annotation pipeline

**DOI:** 10.1101/2020.07.20.211847

**Authors:** Carlos A. Ruiz-Perez, Roth E. Conrad, Konstantinos T. Konstantinidis

## Abstract

**Summary:** High-throughput sequencing has increased the number of microbial genomes from isolates, single cells, and metagenomes available. To analyze and compare these genomes, fast and comprehensive annotation pipelines are needed. Although several approaches exist for genome annotation, these are typically not designed for easy incorporation into analysis pipelines, do not combine results from several annotation databases or offer easy-to-use summaries of metabolic reconstructions in a high-throughput mode. Here, we introduce MicrobeAnnotator, a fully automated pipeline for the comprehensive annotation of microbial genomes that combines results from several reference protein databases to reliably summarize the metabolic potential of the genomes based on KEGG modules.

**Availability:** MicrobeAnnotator is implemented in Python and is freely available under an open source Artistic Licence 2.0 from https://github.com/cruizperez/MicrobeAnnotator

## 1 Introduction

One of the main steps of microbial genome annotation pipelines is the identification and functional prediction of protein-coding genes. Functional prediction is still largely performed manually using searches against reference databases or web-based or command-line tools, e.g., (Seemann, 2014; Tatusova, et al., 2016). These approaches have their own strengths and limitations in terms of speed, database size, and scalability. However, a common theme among them is the difficulty to summarize and compare the resulting large text-based outputs and tables of gene metabolic potential, especially when comparing many genomes.

To fill this gap, we present MicrobeAnnotator, a python-based command-line tool for the automated annotation, summarization and comparison of microbial genomes using multiple reference databases for function prediction. MicrobeAnnotator uses an iterative approach leveraging different bioinformatics tools to search protein sequences against four databases and summarize these annotations into KEGG KO identifiers and modules (Kanehisa, 2019), defined as gene functional units that can be linked to higher metabolic capabilities (pathways), structural complexes, and phenotypic characteristics. It also uses multiple processing cores to simultaneously annotate several genomes or speed up each individual genome annotation. The output of MicrobeAnnotator includes heatmaps and barplots of the metabolic potential of each genome summarizing encoded modules and pathways, and clusters multiple genomes based on metabolic potential similarity.

## 2 Description

### 2.1 Functional gene annotation process

MicrobeAnnotator expects one or more protein sequences in FASTA format as inputs. It implements an iterative annotation pipeline that allows the user to select several search tools and databases including Kofamscan (Aramaki, et al., 2020), Blastp (Camacho, et al., 2009), Diamond (Buchfink, et al., 2015), and Sword (Vaser, et al., 2016), while the databases used include the KEGG Orthology database included in Kofamscan, Uniprot’s Swissprot and Trembl (UniProt, 2019), and NCBI’s RefSeq (O’Leary, et al., 2016). The iterative approach includes five main steps for each set of proteins as follows (Fig. S1): (1) All proteins are searched against the KEGG Ortholog (KO) database using KOfamscan; best matches are selected according to Kofamscan’s adaptive score threshold. (2) Proteins without KO identifiers (or matches) are extracted and searched against Swissprot. This process is repeated twice with searches against (3) RefSeq and (4) Trembl. In all cases the annotation and KO identifiers for each match are saved. Search parameters can also be modified, otherwise the defaults are used (i.e., 40% amino-acid identity, Bitscore 80, alignment length 70%). The last step (5) compiles all annotations and metadata of each match in a table (per genome). KEGG module completeness is calculated for each genome based on the KO identifiers found and a module completeness matrix is produced (and plotted) for all genomes.

**Figure 1.**
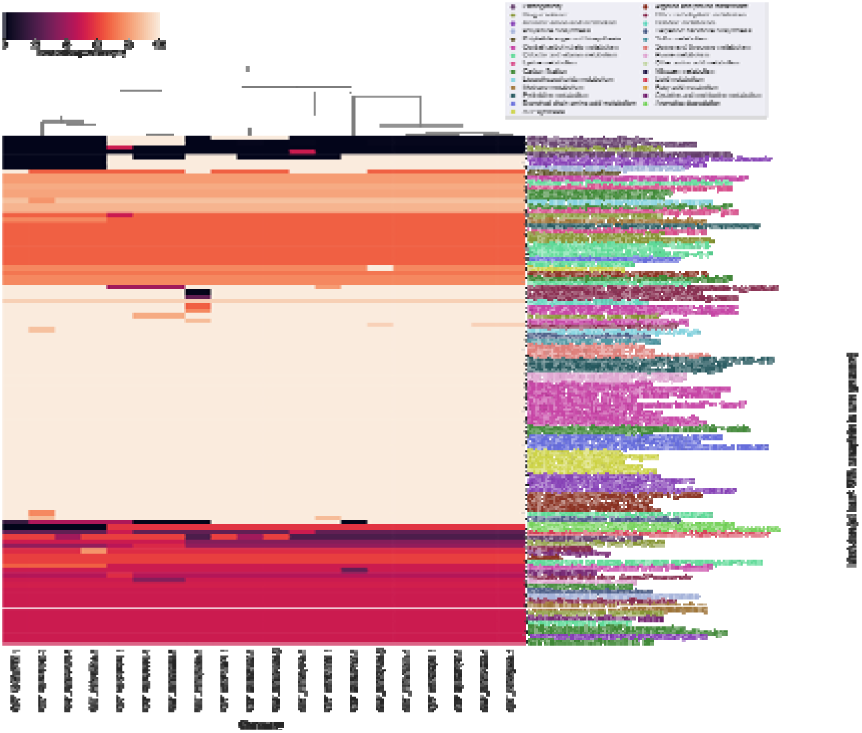
Reduced module completeness (rows) heatmap showing 20 *E. coli* genomes (columns) clustered by metabolic potential.

### 2.2 Output

MicrobeAnnotator results include the compiled annotations and search results for each genome, a summarized matrix of module completeness, and two plots summarizing the metabolic potential of all genomes combined (Figure S3, S4). These outputs are described in the Supplementary Text.

## 3 Results

We compared MicrobeAnnotator with Prokka (Seemann, 2014) and RAST (Brettin, et al., 2015), by annotating a single, randomly chosen *Escherichia coli* genome (GCF_000007445.1). For this comparison, CPU time, number of hypothetical proteins, possibility to run batches of genomes and output formats were assessed. Given that MicrobeAnnotator uses proteins already predicted by other tools, whenever possible all tools started from this step. This was not possible for Prokka and the web version of RAST that perform gene calling by default. The detailed results are shown in table S1. The full version of MicrobeAnnotator had the lowest number of unannotated proteins (4.75%), followed by the light mode with (9.3%), and RASTk (web - 9.8% and local versions 10.5%). As expected, the use of larger databases in the full mode provided a higher number of query proteins with annotations. Notably, 33 out of 330 proteins annotated as hypothetical in NCBI were annotated with MicrobeAnnotator (several related with transposases and phage proteins). Although MicrobeAnnotator has longer run times (396 min full mode and 103 min light mode) compared to Prokka (15 min) and RAST (120 min web and 20 min local), it yields a greater number of annotated proteins and comprehensive summaries (Figure S3, S4). Additionally, MicrobeAnnotator can simultaneously process genomes across threads to significantly speed up the analysis of batch genome collections (details in supplementary section), while the other tools allow the submission of multiple genomes at the same time but they are processed (and reported) independently.

To further highlight the parallelization and output capabilities of MicrobeAnnotator, we performed annotation comparisons of 100 randomly selected *Escherichia coli* genomes. The complete graphical output is found in the Supplementary Text. Genomes and modules are clustered based on module completeness allowing groups of genomes with similar metabolic potential to be identified. For instance, most of the metabolic potential of all *E. coli* genomes is similar with carbon, vitamin, amino acid, and fatty acids metabolism conserved, possibly part of the core genome. Moreover, enterohemorrhagic *E. coli* genomes are separated and clustered together based on the presence of pathogenicity signals (toxins and secretion systems), consistent with previous results (Rasko, et al., 2008). This highlights the ease MicrobeAnnotator offers to quickly annotate and compare groups of genomes. These annotations can be further explored with the annotation files obtained for each genome.

## Acknowledgements

The authors thank Luis Miguel Rodriguez for their comments and suggestions on functionality and documentation of MicrobeAnnotator.

## Funding

This work has been supported by the US National Science Foundation (Award No 1759831)

## Conflict of Interest

none declared.

## Notes

### Competing Interest Statement

The authors have declared no competing interest.

## References

Aramaki, T., et al. KofamKOALA: KEGG Ortholog assignment based on profile HMM and adaptive score threshold. Bioinformatics 2020;36(7):2251–2252.

Brettin, T., et al. RASTtk: a modular and extensible implementation of the RAST algorithm for building custom annotation pipelines and annotating batches of genomes. Sci Rep 2015;5:8365.

Buchfink, B., Xie, C. and Huson, D.H. Fast and sensitive protein alignment using DIAMOND. Nat Methods 2015;12(1):59–60.

Camacho, C., et al. BLAST+: architecture and applications. BMC Bioinformatics 2009;10:421.

Kanehisa, M. Toward understanding the origin and evolution of cellular organisms. Protein Sci 2019;28(11):1947–1951.

O’Leary, N.A., et al. Reference sequence (RefSeq) database at NCBI: current status, taxonomic expansion, and functional annotation. Nucleic Acids Res 2016;44(D1):D733–745.

Rasko, D.A., et al. The pangenome structure of Escherichia coli: comparative genomic analysis of E. coli commensal and pathogenic isolates. J Bacteriol 2008;190(20):6881–6893.

Seemann, T. Prokka: rapid prokaryotic genome annotation. Bioinformatics 2014;30(14):2068–2069.

Tatusova, T., et al. NCBI prokaryotic genome annotation pipeline. Nucleic Acids Res 2016;44(14):6614–6624.

UniProt, C. UniProt: a worldwide hub of protein knowledge. Nucleic Acids Res 2019;47(D1):D506–D515.

Vaser, R., Pavlovic, D. and Sikic, M. SWORD-a highly efficient protein database search. Bioinformatics 2016;32(17):i680–i684.

