## Supplementary Text for "MicrobeAnnotator: a user-friendly, comprehensive microbial genome annotation pipeline"

### 2. Description

MicrobeAnnotator uses an iterative search and annotation approach for the comprehensive annotation of a set of proteins per genome. Figure S1 shows a graphical representation of the MicrobeAnnotator pipeline. The first time a user intends to run MicrobeAnnotator, they will need to build the required databases. For this, the user should run the script MicrobeAnnotator_ DB_Builder, in which it is possible to select the output folders for the databases, the search program to use for future searches (Blast, Diamond or Sword are valid choices), and the number of threads the user wants to run the program with. This script will run four main steps: 1. Download data; 2. Parse annotation data associated with downloaded reference protein sequences; 3. Build SQLite database with annotation data; and 4. Build the protein databases required for the search method selected. If for any reason the program fails at any of these stages, the user can resume the script by selecting the step they want to restart.

Once the databases are built, the program is ready to be used. In this step, the user will provide a set of proteins per given genome (or group of genomes) and will select the search method to use (it is necessary to select the same method as the one used to build the databases). The script MicrobeAnnotator also requires an output directory and the location of the database directory created in the previous step. In addition, the user has the option to customize the annotation by modifying search thresholds (percent identity, bitscore, e-value, or percent alignment across the matching reference protein sequence), and to select whether clustering for the graphical output is desired. Finally, multiprocessing is also allowed with the -t (threads) and -p (processes) options. Each process will annotate a single genome and the number of threads specified will be used in each individual annotation step per genome. For instance, if the user has two genomes and they select -p 2 and -t 4, MicrobeAnnotator will use 8 cores in total for the annotation. It is recommended to not exceed the number of available cores in the system or performance issues will arise.

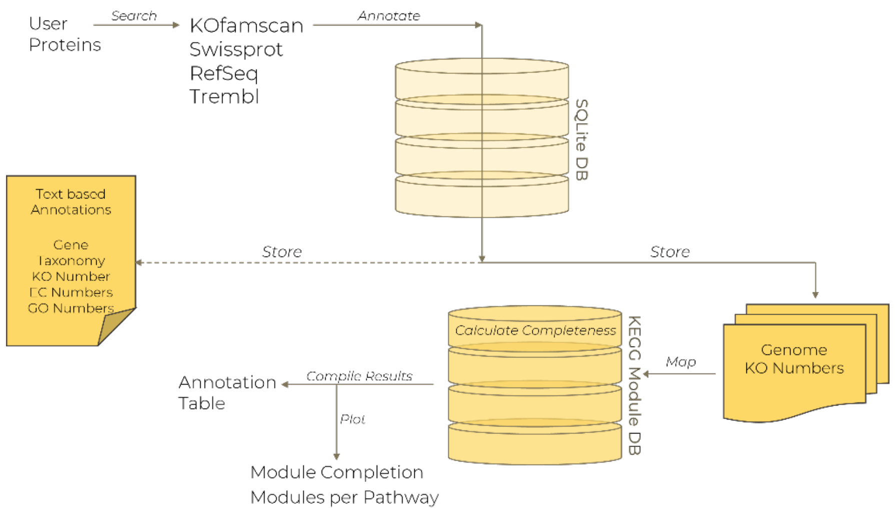


**Figure S1.** Graphical summary of MicrobeAnnotator pipeline. The four main search steps are followed by the annotation of the matches, storing and compiling the results in an easy to parse table and plots.

MicrobeAnnotator will run the pipeline described in Figure S1. Starting from the proteins provided by the user, the program will iteratively search against four main databases (using Kofamscan first and then searching against Swissprot, RefSeq and Trembl, in that order). The original set of proteins is annotated using Kofamscan; the product and KO numbers are stored in a table. Proteins with no KO number associated with them are selected for the next annotation step. This new set of proteins will be searched against Swissprot and the best match for each query is selected. The annotation metadata associated with each matching reference protein, i.e., accession, product, ko number, E.C. number, GO terms and taxonomy, are appended in the annotation table. Again, proteins with no associated KO number are selected for the next annotation step. If the full mode of MicrobeAnnotator is run, the process is repeated for RefSeq and Trembl databases and finally proteins with no matches are annotated as: "Hypothetical protein - no match found", to distinguish between hypothetical proteins in the database (i.e., query proteins matching reference proteins annotated as hypothetical or conserved hypothetical) and no matches, and the process finishes.

The KO number associated with all proteins in each genome (or set of proteins) is extracted and KEGG module completeness is calculated based on the total steps in a module, the proteins (KOs) required for each step and the KOs present in a given genome. KEGG modules are defined as gene functional units that can be linked to higher metabolic capabilities (pathways), structural complexes, and phenotypic characteristics. For example, module M00001 (Glycolysis (Embden-Meyerhof pathway), glucose => pyruvate) is part of the Glycolysis/Gluconeogenesis pathway (00010). These module completeness estimates are compiled in a single table and two plots are generated. The main outputs of MicrobeAnnotator are explained in detail in Table S1.

**Table S1.** Folders and files produced by MicrobeAnnotator

| Result (type) | Description of contents |
|---|---|
| Annotation_results (folder) | Annotations and KO numbers per genome. |
| Kofamscan_results (folder) | Raw and filtered KOfamscan results. |
| Swissprot_results (folder) | Raw and filtered Swissprot results. |
| Refseq_results (folder) | Raw and filtered Refseq results. |
| Trembl_results (folder) | Raw and filtered Trembl results. |
| [prefix].tab (file) | Global table with annotations. |
| [prefix]_heatmap.pdf (file) | Module completeness heatmap. |
| [prefix]_barplot.pdf (file) | Barplot of modules above 80% complete. |

The main heatmap plot shows the module completeness for all modules that are present above 50% complete in at least one genome. The user can select if the program should cluster genomes according to their similarity in module completeness (the same can be done for modules). The second plot is a barplot showing the number of modules above 80% completeness that are associated with a specific pathway. These two plots are explained in greater detail below.

Due to the use of multiple databases for annotation two of which are very large (RefSeq and Trembl) the program may run slow, depending on the computational infrastructure used. With this in mind, we implemented the option for multiprocessing allowing multiple threads to be used in the annotation of a single genome and the option to run multiple genomes simultaneously. We tested the time required to annotate a single *E. coli* genome with different number of threads and repeated this experiment with 50 randomly selected *E. coli* genomes. The system used for this test was part of Georgia Institute of Technology HPC system (PACE). The system had 4 AMD Opteron(tm) 6378 processors with 8 cores per socket at 2400MHz and 2 threads per core (16 threads). The memory available was 30Gb but the maximum used in the test was 11Gb when using 16 threads. Figure S2, shows the improvement in the time required to perform the annotation, which decrease from approximately 100 minutes for a single genome in a single thread to almost half (58 min) using two threads. The speed gains as more computation threads are used continue to increase up to 8 threads, after which the gains are marginal.

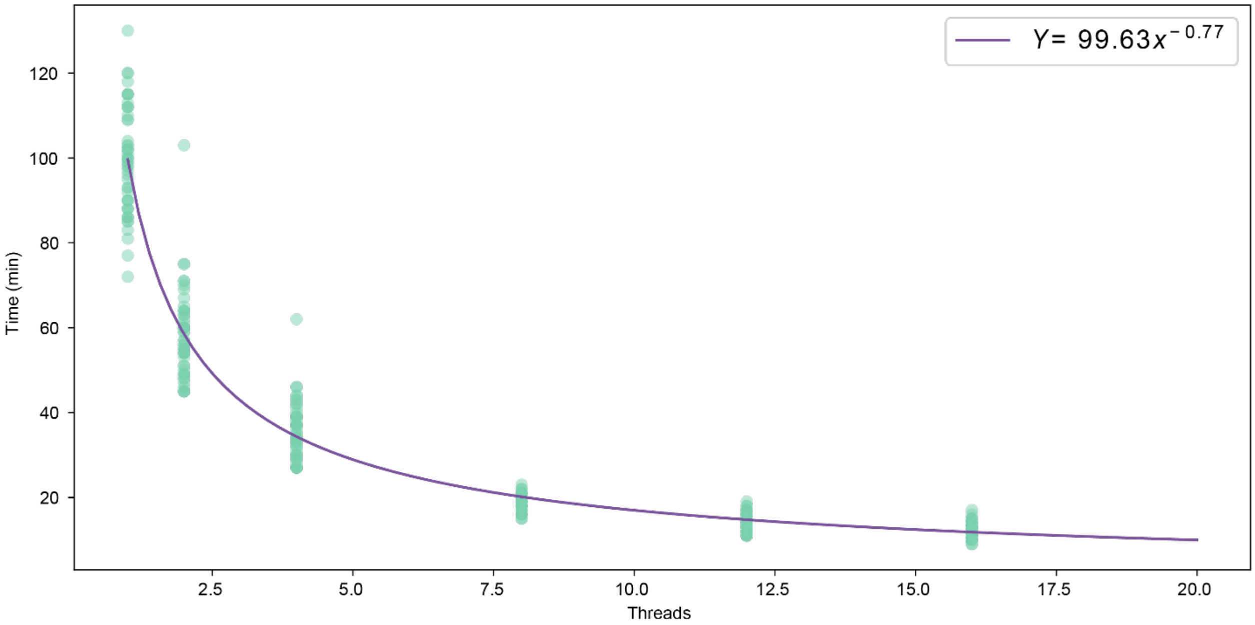


**Figure S2.** Computational time of MicrobeAnnotator when using multiple threads. Note there are significant gains in speed when comparing one vs. eight computation threads per genome. The increase to 12 or 16 threads did not represent major improvements relative to using 8 threads.

### 3. Results

The results of the initial comparison with a single *E. coli* genome (GCF_000007445.1) are shown in Table S1. Both modes of MicrobeAnnotator have the lowest number of proteins without annotations, 220 (4.75%) and 428 (9.24%) for the full and light modes, respectively. These are closely followed by both versions of RASTtk with 506 (9.8%) and 457 (10.5%). Lastly, Prokka annotations resulted in the largest number of proteins without annotation (or annotated as hypothetical). This is also reflected in the time required to complete the annotations, where Prokka is the fastest one at 15 mins compared to 103 and 396 minutes for MicrobeAnnotator (light and full mode, respectively). Depending in the server load and number of jobs, the web version of RASTtk can take up to 120 minutes (this will include queuing and processing time) while the local version will take 20 minutes. One key feature of MicrobeAnnotator is its capacity to simultaneously process multiple genomes and combine their annotation information in a single plot. While the other tools can accept multiple genomes (jobs) at a time, they are processed separately, and their results are individually reported. MicrobeAnnotator on the other hand, will also report the individual annotations of each genome but will also compile a matrix with KO identifiers and report KEGG module completeness in two plots (Figure S3 and S4).

**Table S1.** Comparison of the speed and annotation completeness of MicrobeAnnotator with other tools

| Tool | Proteins | Time (min) | Hypothetical (%) | Batch |
|---|---|---|---|---|
| Original record | 4,634 | NA | 330 (7.12%) | NA |
| MicrobeAnnotator (light) | 4,634 | 103 | **428 (9.24%)** | **Yes** |
| MicrobeAnnotator (full) | 4,634 | 396 | **220 (4.75%)**** | **Yes** |
| RASTtk (web) | 5,187 | 120 | 506 (9.8%) | Yes* |
| RASTtk (local) | 4,634 | 20 | 457 (10.5%) | Yes* |
| Prokka | 4,902 | **15** | 1123 (22.9%) | Yes* |
| Prokka (fast) | 4,902 | **15** | 1192 (24.3%) | Yes* |

* Multiple genomes can be sent for analysis, but they are processed separately.

** Includes hypothetical proteins in database (n=216) and matches not found (n=4).

We tested MicrobeAnnotator with 100 randomly selected *E. coli* genomes from NCBI. All genomes were designated as complete genomes and they could have plasmids, which were included in the computation. For this particular test, we downloaded all proteins from the original accession numbers; alternatively, the user can predict protein-encoding genes from their nucleotide sequences using Prodigal or another program. Figure S3 shows the heatmap generated for the 100 *E. coli* genomes annotated. The color gradient indicates the completeness percentage for each module (y-axis) per genome (x-axis). Figure S4, on the other hand, shows modules above 80% completes grouped based on the pathway they belong to. As expected, the metabolic potential summarized for all *E. coli* genomes is similar in carbon, vitamin, amino acid, and fatty acids metabolism, among others. However, some key differences can be easily identified from the heatmap. For instance, several strains lack the module required for the production of GABA (γ-aminobutyric acid) or the degradation of phenylacetate and homoprotocatechuate (Figure S3, blue boxes). Perhaps more interestingly, depending on the genomes considered, the presence of toxins, secretion systems and antibiotic resistance mechanisms that can help differentiate between different pathogenic strains of *E. coli* such as enterohemorrhagic, enteropathogenic, enterotoxic, among others can be quickly identified (Figure S3, green box).

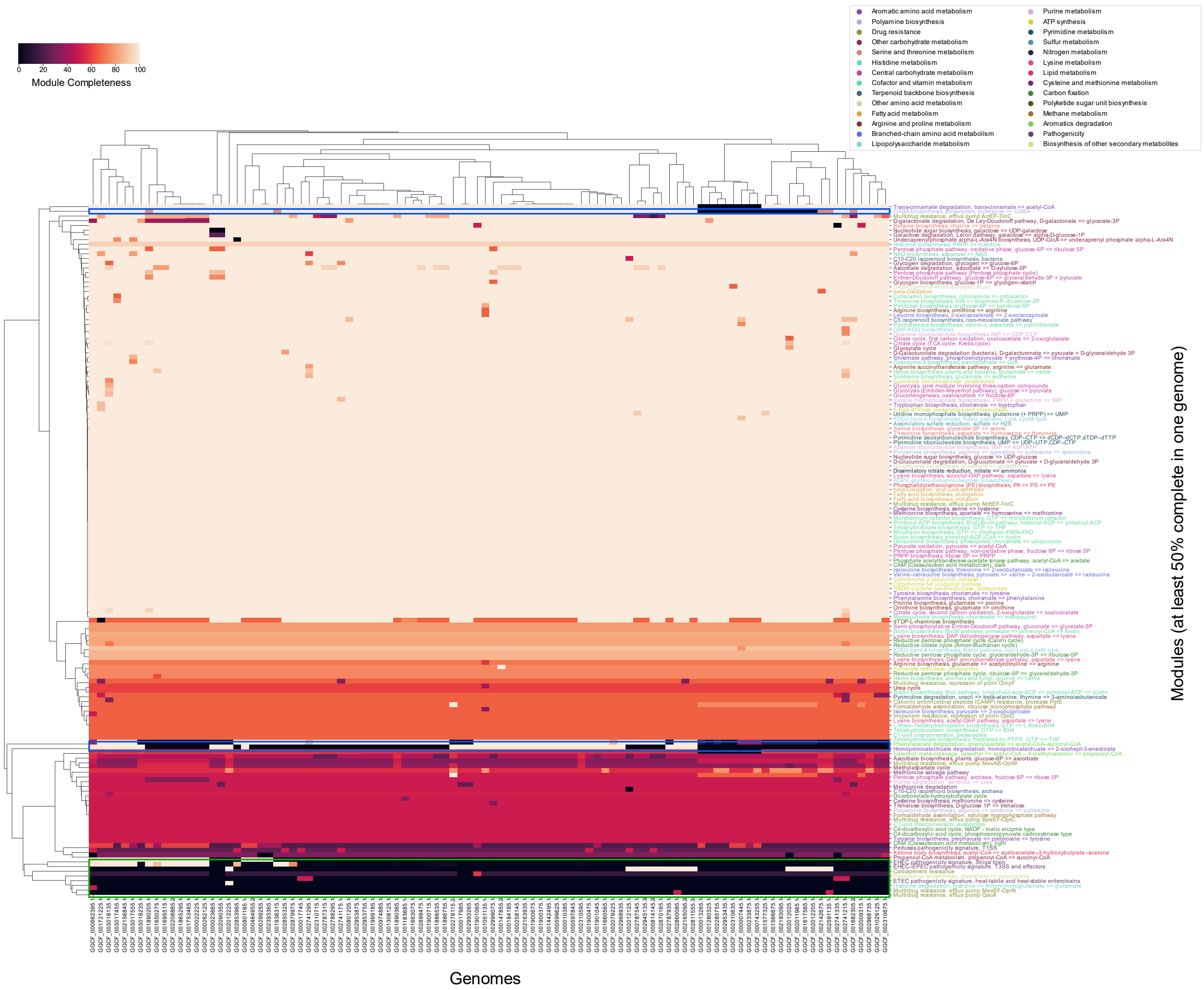


**Figure S3.** Module completeness heatmap of 100 *E. coli* genomes. Note the clustering of genomes based on their overall metabolic potential as well as the clustering of modules that allows an easy interpretation of differentially present modules across genome clusters.

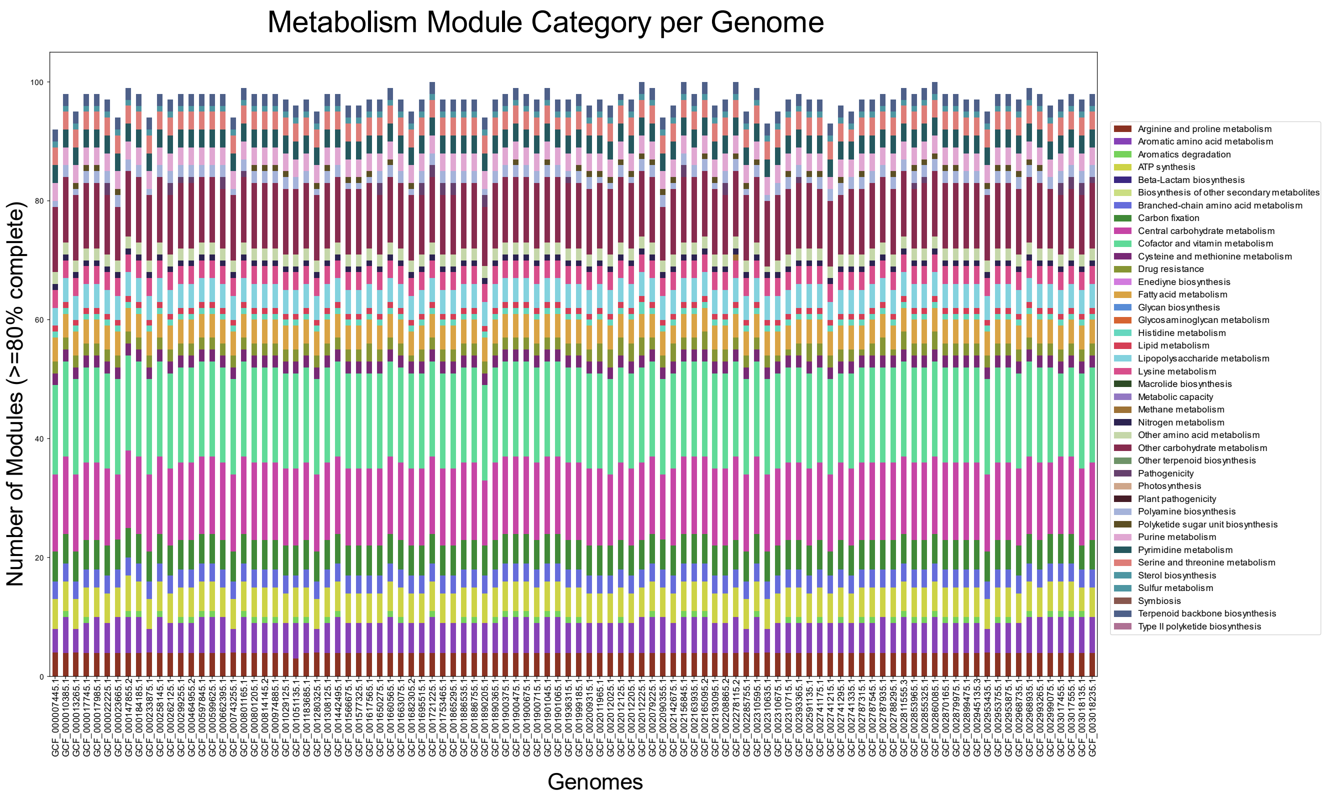


**Figure S4.** Barplots of modules with completeness above 80% grouped by the category (pathway) they belong to. Note the similarity in metabolic pathways exhibited by all *E. coli* genomes analyzed. This plot can also reflect the completeness of genomes, SAGs or MAGs and might reveal missing metabolic pathways in genome reconstructions.
